# Space use fidelity of non-territorial vulturine guineafowl groups is shaped by both environmental and social processes

**DOI:** 10.1101/2025.05.07.652762

**Authors:** Mina Ogino, Brendah Nyaguthii, Danai Papageorgiou, Damien R. Farine

## Abstract

Animals often use consistent areas. Some are territorial, restricting their space use within territorial boundaries, whereas others at not territorial animals but still restrict their space use despite not being constrained by surrounding conspecifics. Staying within a familiar area can provide a range of benefits, such as using previous knowledge (i.e. memory) to efficiently exploit resources or because they can consistently return to key locations (such as a nest or sleeping site). In group-living animals, consistent space use could reduce the complexity of decision-making time (e.g. by choosing among known foraging sites), facilitating group cohesion. However, to date, little research has explicitly asked what factors determine whether groups use consistent areas. Here we used repeated movements by groups of vulturine guineafowl (*Acryllium vulturinum*)—leaving and returning back to the same areas in response to seasonal conditions—to examine and disentangle social processes from spatial and ecological factors that might shape the distribution of animals over space. Specifically, we quantified (i) how groups distribute themselves over the landscape, (ii) if their space use is consistent across seasons with similar environmental conditions, (iii) how different social and spatial factors shape the consistency of space use by groups over time, and (iv) how social and spatial factors affect home range overlap between groups. We found that groups were highly consistent in their space use over time and that home ranges were distinct across groups. Fidelity to the core home range area was higher when group composition was more stable, while overall home range fidelity was higher when groups recently experienced milder ecological conditions. Overlap in core areas and the overall home ranges among groups were greater among groups that shared roosts and groups that were fused in the previous season. Home range overlap was also lowest during long intermediate seasons (i.e. a sampling period that immediately follows intermediate season conditions, as opposed to sampling periods that followed dry or wet conditions), suggesting that extended intermediate conditions allow groups to increasingly partition their overall space use. These results provide insights into how the movement decisions by groups, the distribution of animals, and group-level space use emerge, and the role of social and ecological conditions as potential precursors to territoriality.

## Introduction

Animals often consistently use the same areas over time to live and reproduce (so-called home ranges (Burt, 1943)). For example, migratory birds can come back to the same breeding territories or use the same wintering sites consistently across years (Grist et al., 2014; Rohwer & Anderson, 1988; Somershoe et al., 2009). The most extreme cases of consistent space use are typically territorial animals that establish territories to secure exclusive access to resources and defend their territories from competitors (Wilson, 2000). Space use by territorial animals is therefore highly constrained by territorial boundaries. Here, neighbouring territory holders can express either tolerance or escalated aggression towards each other depending on the underlying social contexts, adding another social layer shaping space use (Christensen & Radford, 2018). On the other hand, many species do not establish territories (i.e. non-territorial animals), and thus their space use is not restricted by conspecifics defending a given area. Non-territorial animals are therefore able to choose to stay or leave an area. Yet, even in non-territorial species we often observe that individuals express consistent space use, for example, by establishing home ranges (Burt, 1943) or by repeatedly coming back to the key locations (e.g. (Andersson, 1978, 1981)).

Several factors are known to facilitate the emergence of consistent space use (i.e. high fidelity in space use) by animals [reviewed in (Piper, 2011)]. One key factor is the use of memory (Fagan et al., 2013; Ranc et al., 2021; Riotte-Lambert et al., 2015). Individuals can remember where they foraged before and can repeatedly come back to the area that gave them the best foraging outcome. By repeatedly using the same area, they can also gain familiarity with the surrounding habitat configuration (Piper, 2011) which allows them to use energetically efficient paths (Gallagher et al., 2017; Shepard et al., 2013), to avoid areas of high predation risk or efficiently evade from predators (landscape of fear (Gallagher et al., 2017; Gaynor et al., 2019) also (Clarke et al., 1993; Yoder et al., 2004)), and to predict the distribution of resources over time (learning phenology or estimating renewal rates) (Brown et al., 2008). Consistent space use can also emerge from the need to return to key locations (i.e. central-place foraging). For example, breeding seabirds need to travel between foraging sites and their nesting colony to provision their offspring, thereby limiting their space use according to the travel distance from their nesting site (Elliott et al., 2009). African elephants *Loxodonta africana* are also known to repeatedly use the same sinkholes across years (Fishlock et al., 2016). As well as providing an essential resource (water), these sinkholes act as the key locations to facilitate social interactions among individuals. Thus, consistent space use can emerge from both limitations to movement as well as to enable animals to efficiently exploit resources.

Space use is also governed by interactions that animals have with others, including both social partners and competitors. Space use by individuals is often tied to the presence of social associates in the area. For example, golden-crowned sparrows (*Zonotrichia atricapilla*) consistently use the same wintering sites across years, if they find their social partners from the previous winter (Madsen et al., 2023). Individuals can also preferentially use different spaces from other individuals to avoid interference or indirect competition. Competition can result in individuals being excluded from an area or result in them arriving at a resource that has already been depleted. The ideal free distribution hypothesis (Fretwell & Calver, 1969) proposes that—as a result of competition—individuals should evenly distribute themselves according to how resources are distributed over space. The level of competition can also vary corresponding to the ecological condition. When the environmental conditions are milder and the resource is abundant, the competition may be relaxed, allowing individuals to have greater overlap in their space use. On the other hand, when the environmental condition is harsher and the resource becomes scarce, the competition over resources becomes severe, driving greater partitioning of resources (and thus space use).

The dynamics of competition that drive space use patterns among individuals should equally affect those by groups. However, in species where individuals preferentially interact among themselves and move as cohesive social units (Davis et al., 2022), group members face a number of additional challenges that can increase the benefits of maintaining high space use fidelity (i.e. a consistent home range). For example, groups need to maintain cohesion and reach a consensus on where to go despite potential conflicts of interest among group members (Conradt, 2012). Solving social challenges could be linked to consistent space use. For example, when space use by groups is more predictable (i.e. restricted to a more limited set of options), then groups may need a shorter time to reach a consensus about where the group is going next (because of the limited number of options, see (Tanner & Hemingway, 2025) at the individual level). Similarly, they may be more efficient in moving through familiar space because they share a common end-goal and knowledge about how to get to the destination. Previous work has shown that collective movements are slower than individual movements, likely as a result of the constant process of decision-making, which can impose both energetic (Harel et al., 2021; Klarevas-Irby et al., 2025) and time costs. Having shared spatial knowledge among group members could reduce the need to engage in constant decision-making actions as groups navigate their way through the landscape. Further, group foraging can lead to greater intake rates when resources are depletable (i.e. when there is more competition), because groups can better sense the environment (Clark & Mangel, 1984; van der Post & Semmann, 2011) and because foraging as a group reduces the chance of selecting a patch that is depleted (Beauchamp & Ruxton, 2005). Thus, consistent space use can facilitate cohesion and foraging in a group can be beneficial when exploiting a consistent home range.

The ability for groups (or individuals) to express consistent space use is also subject to social change over time. First, demographic changes due to death, birth, and dispersal will change both the distribution of competitors (thus freeing up resources) and the membership of social groups (Ekman et al., 1981; Krebs, 1971). Since different individuals have different needs or experiences (reviewed in (Ogino et al., 2023)), fidelity in space by groups may become lower when new individuals join the group or when some individuals (correspondingly, some information) are lost from the group ((Galef Jr & Whiskin, 1997; Stanley et al., 2008)). Second, in a number of animal species, individuals and groups preferentially interact with or avoid certain conspecifics or sets of groups (Christensen & Radford, 2018; Mirville et al., 2020). One extreme example is multilevel societies, where the decisions to merge and split are made by groups (rather than individuals) and where sets of groups can move as one cohesive supergroup (Grueter et al., 2020; Papageorgiou & Farine, 2021). In such societies, previous history between groups can also determine what knowledge the group may possess and the tendency of groups to come back to the area where other groups can be found. Thus, social stability and/or previous group-level events are likely to play important roles in determining fidelity in space use.

To date, only a few studies (e.g. (Hendrix et al., 2024)) have investigated the relative roles of social and spatial factors in shaping the consistency of space use by groups. This is because disentangling social processes from spatial processes is impossible when individuals use different areas from each other (i.e. individuals or groups with different home ranges also experience different physical and ecological environments). One way to address this question is by studying societies where groups are not territorial and have highly overlapping home ranges, as this allows us to partition social effects from ecological drivers in shaping space use. Species with spatially overlapping but non-territorial home ranges—like those found in multilevel societies—thus provide the ideal opportunity to disentangle social from spatial factors driving consistent space use.

We study a population of vulturine guineafowl (*Acryllium vulturinum*) that consists (on average) of 23 non-territorial groups (15-65 individuals per group (Papageorgiou et al., 2019)). In savanna in East Africa, there are two extreme seasons, wet and dry seasons, which are interspersed by intermediate seasons. Both wet and intermediate seasons are favourable, and vulturine guineafowls opportunistically breed during wet seasons. Group membership is typically stable over time. Different groups tend to merge and form supergroups under harsher ecological conditions (fission and fusion events during dry seasons and droughts, i.e. forming a multilevel society), but groups almost always split back to the original group compositions when the ecological conditions get more favourable (intermediate seasons; and then can split into breeding subgroups during wet seasons (Nyaguthii et al., 2025)). Groups have been reported to shift and expand home ranges during dry seasons (Papageorgiou et al., 2021), with groups returning to preferred home ranges (and home ranges contracting again) during more favourable conditions (intermediate and wet seasons). Finally, groups are completely non-territorial and often co-exist spatiotemporally (both during the day and at communal night-time roosts (Papageorgiou, Cherono, et al., 2024)). Together, these features mean that we can observe groups returning to the same home range after experiencing demographic change and varying amounts of harshness, while also obtaining data from replicated groups sharing the same (or similar at least) ecological and physical environmental conditions.

We quantified space use dynamics in non-territorial vulturine guineafowl groups by combining long-term GPS data with group membership data inferred from the long-term census collected from the study population. Specifically, we investigated (i) how non-territorial groups distribute themselves over space, (ii) whether space use by groups is consistent across seasons with similar environmental conditions, (iii) what social processes (changes in within-group and between-group social environments) and spatial factors impact on stability in space use by groups over time, and (iv) if social and spatial processes shape the overlap in space use among groups. We quantified space use using the utilisation density (UD) and with core area (50% flat polygons) determined from the UD. The 50% flat polygons represent the core area that groups used the most frequently. Groups may utilise different parts of their home ranges under different contexts (e.g. foraging, resting, and so on), and thus, space use of these different parts might be shaped by different factors. From the field observation, we noted that home range fidelity of vulturine guineafowl groups is largely consistent over time, but that home ranges and the extent of home range overlaps between groups can vary. We predict that this variation is likely to be linked to social processes (especially changes in group membership) and environmental conditions (especially when groups experienced harsh conditions between two similar seasons). Doing so allows us to disentangle the relative role of social from spatial processes governing the distribution of animals over space.

## Methods

### Data collection

#### Study system and study period

Vulturine guineafowl are endemic to the East Africa. We studied a population of vulturine guineafowl in the savannah area at the Mpala Research Centre in Kenya. The population are usually observed within a small area (approximately 8 km^2^) around the research centre under mild ecological conditions. In harsh conditions, the population will move outside of this area in search of resources but immediately return to the area around the research centre when the conditions become milder (following rainfall events) (Papageorgiou et al., 2021), indicating that this is a preferred area for these groups to be in.

The majority of individuals in the study population were colour-banded for individual identification (approximately 90 % of individuals in the population, 96–698 individuals with rings, varying over time depending on environmental conditions, dispersal and recent breeding events). Based on the colour bands, we collected census data almost every morning (6:00) and evening (18:00) since 2016. We drove throughout the study area to find groups of vulturine guineafowls to record the colour bands of individuals, the location where the individuals were observed, time, date, the total number of individuals observed, whether the exact number of individuals was recorded, and whether all ringed individuals were recorded. We tried to ensure that all groups were regularly observed throughout the study period. The study population consisted of 12-22 stable groups (see the following section for the detailed detection methods) during intermediate seasons (these stable groups merge and form supergroups under harsher ecological conditions). Each group had at least one GPS-tagged individual (we aim to tag 10% of individuals in each group). In this study, we used GPS data points collected every 5th minute during the daytime to estimate ranging behaviour.

#### Classifying season types

We classified days between June 30^th^ 2018 and October 15^th^ 2023 into four seasonal conditions (wet, intermediate, dry, and drought) based on normalized difference vegetation index (NDVI) and rainfall. NDVI was determined following the previous study in the same study population (Papageorgiou et al., 2021). Specifically, the start of a wet season was defined as the day marking a large rainfall event (typically approximately 20 mm in a day) and followed by an immediate increase in NDVI to be above 0.45 (i.e. the rainfall causes grasses to become green). Typically, NDVI drops towards the end of the wet season. We defined the approximate date when NDVI becomes less than 0.45 as the start of an intermediate season (i.e. rainfall cannot maintain the greenness of grasses). As NDVI further drops throughout an intermediate season, we defined the period when NDVI dropped below 0.25 as a dry season. Finally, when NDVI became lower than 0.2, we classified such a period as drought. Our study period included 5 wet, 6 intermediate, 2 dry seasons, and 2 droughts when using this classification system. These cut-offs for NDVI values were largely determined by the field experience and the observed seasonal changes in the broader ecosystem (e.g. changes in vegetation and the behaviour of other species).

For the purpose of our study, we focused only on the space use during intermediate seasons. Intermediate seasons represented the period when the space use of groups was not restricted by nesting sites or driven by harsh ecological conditions, such as the availability of water, while groups seemed to preferentially return to the area from the observation. However, because intermediate seasons varied in length, we also divided long seasons into separate sampling periods, thus allowing us to make each sampling period approximately the same length (1.5 to 2.5 months) to make home range estimations comparable (see the section below, ‘*Estimating how groups used space*’ for the details for home range estimations). For example, if an intermediate season lasted longer than 3 months, we divided this season into two sampling periods, and estimated home ranges for each sampling period. This resulted in 9 wet, 12 intermediate, 4 dry, 7 drought sampling periods. While we contrasted between sampling periods (including in ‘consecutive’ intermediate seasons), hereafter we refer to sampling periods as distinct ‘seasons’ for ease of clarity.

#### Identifying group membership and estimating group sizes

We determined group membership for each sampling period following the methods established in previous studies on the same system (Ogino et al., 2023). Inferred group membership closely matched the general patterns of group membership tracked by the field team. When males (the philopatric sex) from a detected group during the focal sampling period were detected with males from other groups in previous sampling periods, we defined the group as experiencing fission-fusion events. We assigned a new ‘moving unit ID’ to identify and keep track of which sets of groups in the most recent intermediate seasons previously merged to form a supergroup (typically during drier seasons) and expressed collective behaviour and when the supergroups split back to the original groups (during wetter seasons).

### Data processing

We collated five possible predictors of home range fidelity (Figure 1). Our analyses focused on the fidelity of space use from one intermediate season to a subsequent intermediate season. This is because intermediate seasons appear to capture periods when groups maintain their preferred home range (as opposed to dry seasons when they make large movements in search of resources) and because home ranges are not affected by reproductive behaviours (during wet seasons) (Papageorgiou & Farine, 2020).

**Figure 1.**
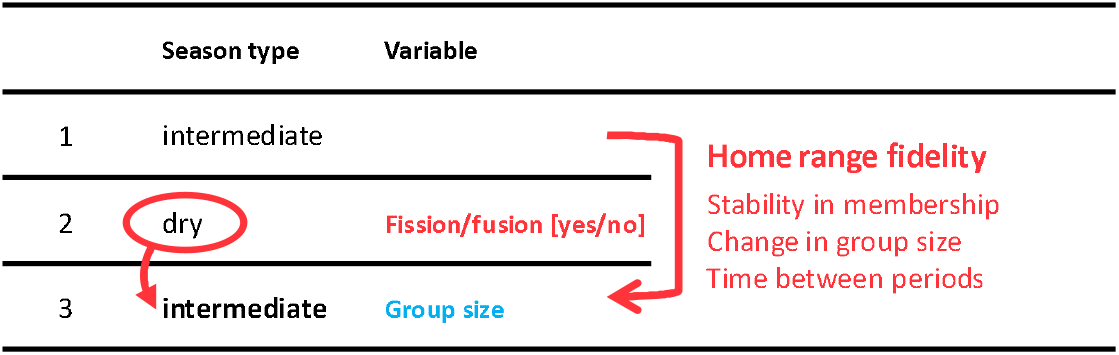
Potential social and ecological drivers of space use fidelity across seasons by vulturine guineafowl groups. We determined home range fidelity, comparing home ranges estimated from one intermediate season (e.g. the previous intermediate season, sampling period 1) to the next (the focal intermediate season, e.g. sampling period 3). We determined the ecological condition prior to the focal season (e.g. a dry season, sampling period 2, shown by the arrow pointing to the red circle). We also determined social factors during the focal season (e.g. group size) and the dynamics between intermediate seasons. The latter includes group-level fission-fusion events that may have occurred between the previous intermediate season and the focal season (i.e. between sampling periods 1 and 3), and changes in group size and group membership since the previous intermediate season. We fit these variables as predictors and home range fidelity values as response variables in GLMMs.

#### The season types right before the focal intermediate seasons

To test whether ecological conditions shape the subsequent space use, we determined the ecological condition prior to the focal season (Figure 1).

#### The number of days between the previous and focal intermediate seasons

As a proxy for memory use, we determined the number of days between focal sampling periods (Figure 1). We subtracted the end date of the previous intermediate season from the start date of the next intermediate season.

#### Stability of group membership

To test whether changes in group membership affect the patterns of space use by groups, we determined how similar the members in moving unit IDs were from one focal sampling period to another (Figure 1). We quantified the similarity in group membership (i.e. which individuals moved together with) using the Jaccard similarity index of the membership (herein, *Jaccard similarity of the membership*). The Jaccard similarity was determined by dividing the number of common individuals by the total number of detected individuals in both focal sampling periods. In addition to this, by tracking the inferred group IDs, we determined if the focal group merged with or split from other groups between focal sampling periods (for home range fidelity) and if the focal groups split from the same parent group after the previous sampling period (for home range overlaps).

#### Group size

Identifying group sizes is not straightforward. First, not all birds are banded, and thus, the true group size might not be captured by the number of individuals identified as a part of each group. Second, accurate census data can sometimes be hard to obtain (e.g. when a group is moving through dense vegetation we can miss some individuals). Thus, it is not always unambiguous to know exactly how many unmarked individuals are within each group. What we do to address this is to obtain a count of the number of individuals in total (both marked and unmarked) we think are in a group that we observe. This then generates multiple instances of these counts—which is complicated by the fact that groups are sometimes joined with other groups. To generate an accurate estimate of the size of each moving unit in each sampling period (i.e. groups in intermediate seasons), we extracted the corresponding census observations that included males from a given moving unit ID and calculated the median number of unbanded individuals observed together across each of these observations. We used the median number of unbanded individuals detected as a part of the group for each census observation as the moving unit (group) sizes in the following analysis.

#### Estimating how groups used space

Using 5-minute GPS data, we estimated home ranges and core areas of all GPS-tagged adult males (the philopatric sex) for each sampling period. We used 95% autocorrelated kernel density estimation (AKDE) in the *ctmm* package (Calabrese et al., 2016; Fleming, 2022) in R (Team, 2021), to estimate the UD of each group (herein, *overall home range*). From each UD, we also determined the core area where one can find the focal group with a 50% probability as flat polygons (herein, *50% flat polygon*), using the *ctmm* R package (Calabrese et al., 2016; Fleming, 2022). The 50% flat polygons represent the core area that groups used the most frequently potentially for key activities such as foraging and resting.

#### Quantifying consistency and overlap in space use

To investigate if space uses by groups were consistent across time (i.e. across sampling periods) and how groups distribute themselves over the landscape, we defined fidelity in space use as within-group home range overlap between intermediate seasons, and between-group overlap as home range overlap among groups within an intermediate season. We calculated overlap for all pairs of seasons within groups (see Figure 1 for an example) and for all possible combinations of groups using overlap function (Winner et al., 2018) in *ctmm* package (Calabrese et al., 2016; Fleming, 2022). If groups were using the whole available area without any preference, then we expect them to have either (1) lower home range fidelity and higher home range overlap or (2) no difference between home range fidelity and home range overlap. We thus ran Kruskal-Wallis multiple comparisons (p-value: Bonferroni method) and Dunn’s post-hoc test in the FSA package (Ogle, 2025) to determine if overlap values in different categories (fidelity in UD and 50% flat polygon; overlap in UD and 50% flat polygons among groups) were significantly different from one another.

### Data analysis

#### Testing how within-group stability in space use is impacted by social and environmental processes

To investigate how within-group stability in space use is impacted by social processes, we built generalized linear mixed models (GLMMs; beta family) using the *glmmTMB* package (Brooks et al., 2017). We fit home range fidelity (home range overlap within groups under the same season types in the different sampling periods) as a response variable. As fixed effects, we fit the Jaccard similarity of the membership, change in group size since the previous intermediate season (scaled absolute values), whether the group merged with or split from other groups since the previous intermediate season [binary], group size in the focal intermediate season (scaled), the season types right before focal intermediate season [wet, intermediate, dry], and the number of days between the previous and focal intermediate season (scaled). The *inferred stable group ID* was fitted as a random effect. We fit the same models with the UD and 50% flat polygon as response variables. We visually confirmed the model fit using the DHARMa package (Hartig, 2022).

#### Testing how space use overlap among groups is shaped by social and ecological processes

We built GLMMs (beta family; *glmmTMB* package) to test whether spatial and social processes between groups shape overlap in space use among groups. We fit home range overlap between each pair of groups (during the same sampling period) as a response variable. We then fit the following variables as predictors: the similarity among the sets of roosting sites groups used during the focal intermediate season (*cosine similarity* in roosting site use), the size of the largest of the two groups being compared, whether the two groups split from the same parent group after the previous intermediate season [binomial], the size difference between the two groups, and the season type immediately prior to the focal intermediate season [wet, intermediate, dry season]. We also included a unique identifier for each pair of groups (i.e. ‘inferred group ID 1_inferred group ID 2’) as a random effect. We fit the same models for the overlap determined from 50% flat polygons. No critical dispersion was observed in any models and the model fit was visually inspected using the DHARMa package (Hartig, 2022).

## Results

### Space use by groups is consistent over time despite overlap with other groups

Groups were generally consistent in their overall home range from one intermediate season to the next (Figure 2a). During a given intermediate season, group home ranges also overlapped with each other to a high degree (Figure 2b). Overall, in a given intermediate season the home range fidelity (from the previous intermediate season) was significantly greater than the home range overlap between groups (Z=-16.722, P<0.001). Both home range fidelity and between-group home range overlap were highly variable (Figure 2).

**Figure 2.**
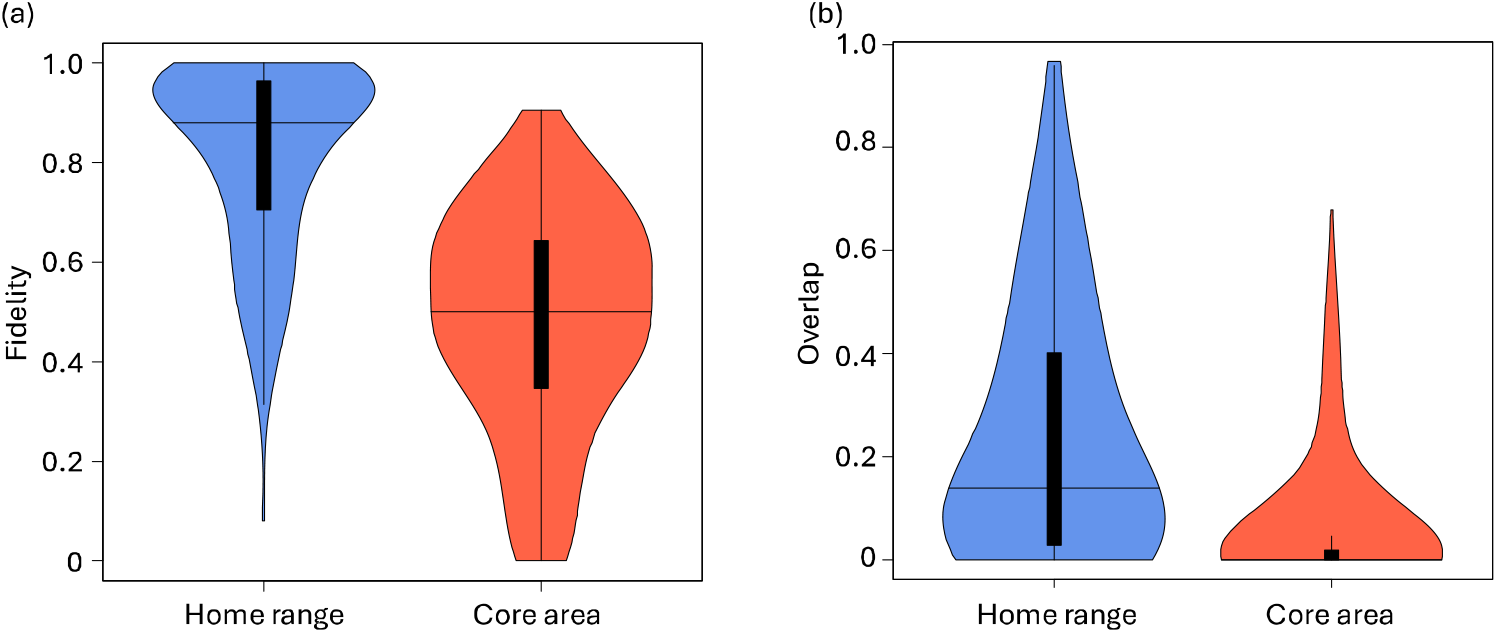
Fidelity in group home range and between-group home range overlap. (a) shows home range (blue) and core area (red) fidelity within groups, and (b) represents home range (blue) and core area (red) overlap between groups. The middle horizontal line in each box shows the median values, and the upper and lower ends of the thick black line show the upper and lower quantiles of the distribution.

Groups were also consistent in their use of core areas (Figure 2), but their fidelity in their core area use was substantially lower than fidelity in their overall home range (Z=-6.482, P<0.001). Groups also overlapped in their core areas, but these were more distinct than the broader home ranges (Z=-10.773, P<0.001; Figure 2). Overall, core area fidelity was significantly higher than core area overlap between groups (Z=-22.605, P<0.001, Figure 2). Both core area fidelity and overlap were highly variable (Figure 2).

### Social and ecological processes predict temporal variation in home range fidelity

Both social processes and the ecological conditions that groups experienced played a role in shaping the variation in the fidelity of the overall home range from one intermediate season to the next (Table 1). Larger groups showed higher fidelity in their overall home ranges, but stability in group membership, changes in group size, and recent group-level fission-fusion events did not affect the fidelity of the overall home range. The overall home range fidelity of groups was lower between intermediate seasons that were separated by longer periods of time. The type of condition experienced between intermediate seasons also mattered. Home range fidelity tended to be higher right after groups experienced wetter ecological conditions than after groups experienced harsher ecological conditions (dry seasons).

**Table 1.**
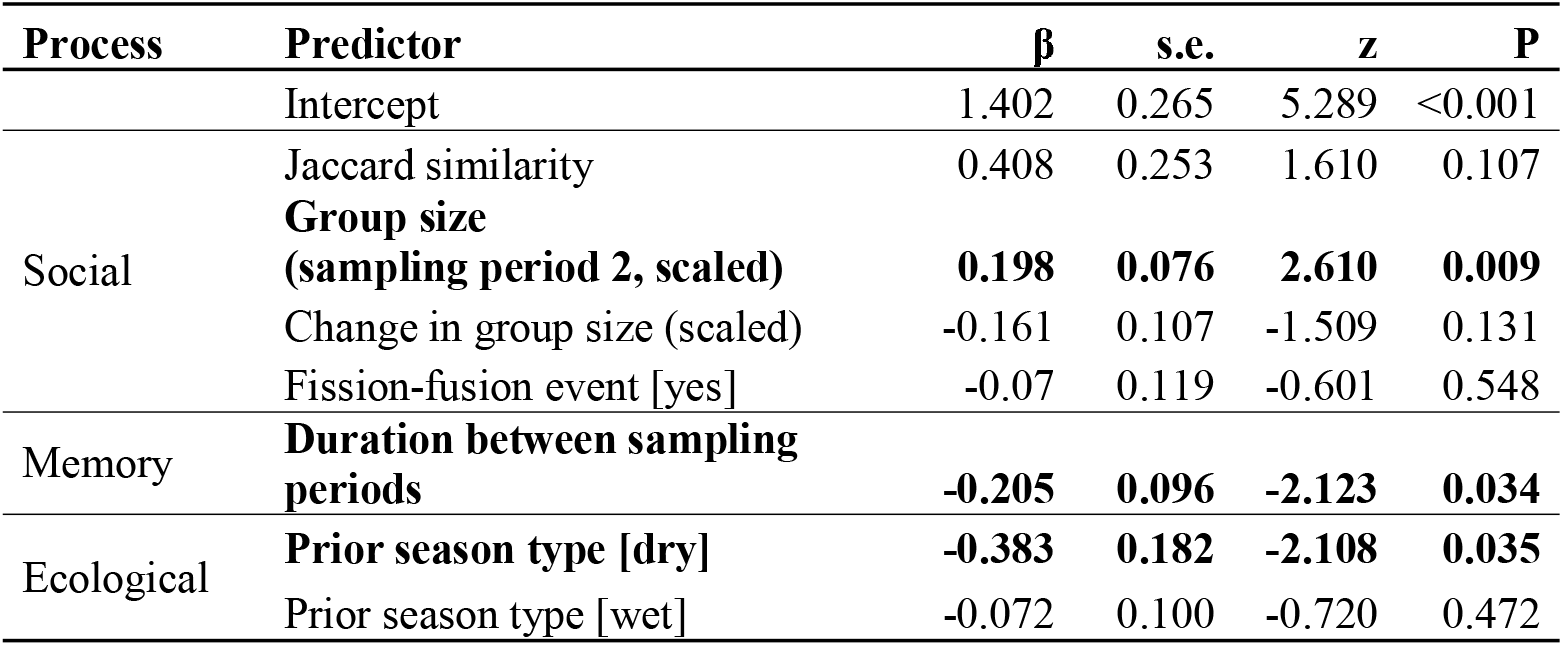
The role of social factors, ecological processes, and memory in driving variation in overall home range fidelity. Results of a GLMM (beta family) with fidelity in UD as a response variable. A total of 17 groups across 11 sampling periods were included in the analysis. Group ID was fitted as a random effect (variance = 0.142; SD = 0.376).

The patterns for core area use were mostly similar (Table 2) but with some key differences. Larger groups had a higher fidelity in their core areas. Groups with more stable membership (higher Jaccard similarity) showed higher fidelity in their core ranges. Likewise, groups that changed size had lower fidelity in core areas. The occurrence of recent group-level fission-fusion events did not affect the fidelity in core areas. The duration of time between intermediate seasons did not affect fidelity in core area use, but fidelity in core areas tended to be higher after groups experienced wetter ecological conditions relative to harsher ecological conditions.

**Table 2.**
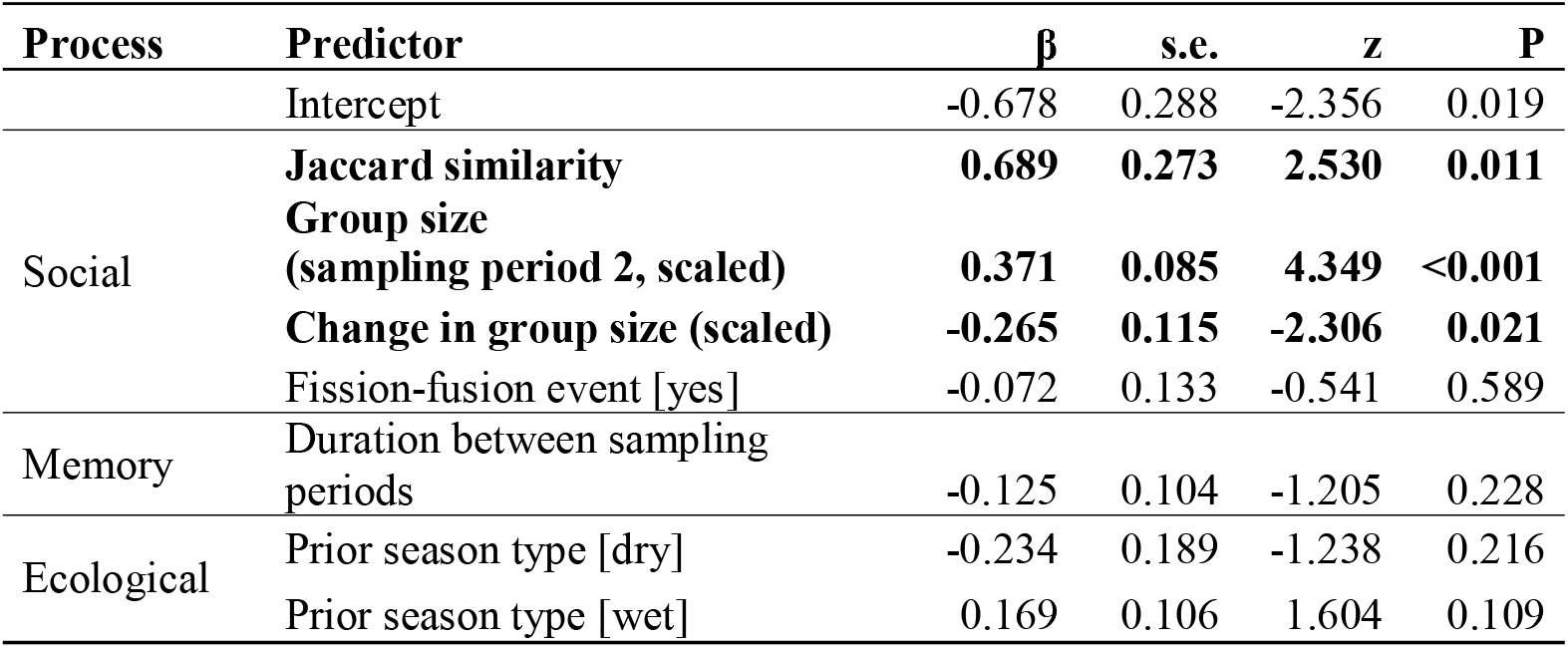
The role of social factors, ecological processes, and memory in driving variation in fidelity in the core area used by groups (50% flat polygon). Results of a GLMM (beta family) with fidelity in the core area used as a response variable. Group ID was fitted as a random effect. A total of 17 groups across 11 sampling periods were included in the analysis. Group ID was fitted as a random effect (variance = 0.224; SD = 0.473).

### Social and ecological processes predict variation in home range overlap among groups

Social and ecological processes also affected the overlap in the overall home range among groups (Table 3). Home range overlap was higher among groups that used more similar sets of roosting sites, but was smaller when the size difference between groups was larger. Groups that previously merged and split (fission-fusion event) had higher home range overlap than sets of groups that were not previously merged. The size of the larger group did not affect the amount of home range overlap among groups. Home range overlap was also significantly lower during sampling periods that followed a previous intermediate season sampling period (i.e. within a long intermediate season), with groups also having a lower overall home range overlap following wetter ecological conditions than after harsher ecological conditions (dry seasons).

**Table 3.** The role of social and ecological processes in driving variation in between-group overlap in overall home ranges. Results of a GLMM (beta family) with the between-group overlap in UD used as a response variable. The combination of compared group IDs was fitted as a random effect (variance = 0.142; SD = 0.376). A total of 17 groups (resulting in 188 combinations of groups) across 11 sampling periods were included in the analysis.

| Process | Predictor | $\beta$ | s.e. | z | P |
| --- | --- | --- | --- | --- | --- |
|  | Intercept | -1.893 | 0.072 | -26.35 | <0.001 |
| Social | **Cosine similarity of roost sites** | **2.492** | **0.059** | **42.57** | **<0.001** |
|  | **Fission-fusion event [yes]** | **0.340** | **0.078** | **4.34** | **<0.001** |
|  | **Differences in group sizes (scaled)** | **-0.030** | **0.015** | **-1.97** | **0.049** |
|  | larger group size (scaled) | -0.028 | 0.027 | -1.02 | 0.309 |
| Ecological | **Prior season type [dry]** | **0.687** | **0.072** | **9.52** | **<0.001** |
|  | **Prior season type [wet]** | **0.228** | **0.030** | **7.56** | **<0.001** |

Relatively similar patterns were found for the overlap in core areas among groups but with some key differences (Table 4). Groups that used more similar sets of roosting sites had higher core range overlap, whereas groups with very different sizes had less overlap in core ranges. Groups that recently merged and split (fission-fusion event) had more overlapping core ranges, but the effects of group size differed from the overall home range overlap. Specifically, the size of the larger group was more important than the difference in group sizes in determining core range overlap. Core area overlap was significantly higher right after groups experienced harsher ecological conditions (dry seasons), with the opposite (but not significant) pattern emerging after wetter ecological conditions (unlike the overall home range).

**Table 4.**
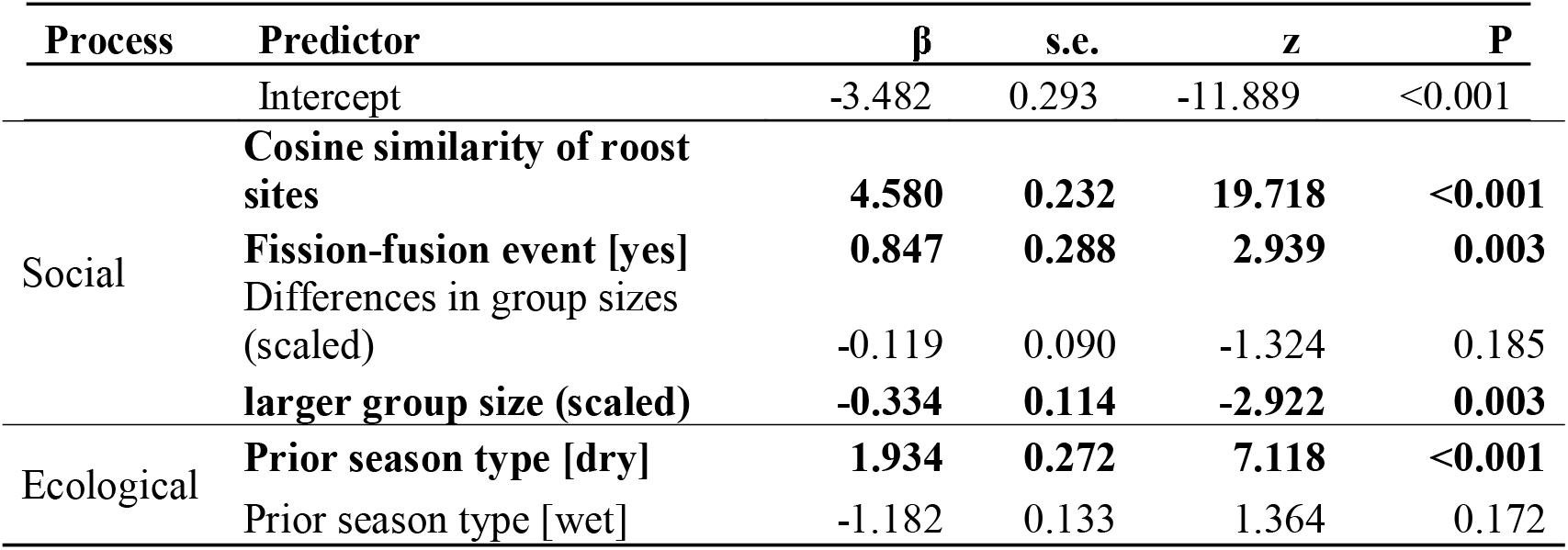
The role of social and ecological processes in driving variation in between-group overlap in core home range areas. Results of a GLMM (beta family) with core home range overlap between groups used as a response variable. The combination of compared group IDs was fitted as a random effect (variance = 6.772; SD = 2.602). A total of 17 groups (resulting in 188 combinations of groups) across 11 sampling periods were included in the analysis.

| Process | Predictor | $\beta$ | s.e. | z | P |
| --- | --- | --- | --- | --- | --- |
|  | Intercept | -3.482 | 0.293 | -11.889 | <0.001 |
| Social | **Cosine similarity of roost sites** | **4.580** | **0.232** | **19.718** | **<0.001** |
|  | **Fission-fusion event [yes]** | **0.847** | **0.288** | **2.939** | **0.003** |
|  | Differences in group sizes (scaled) | -0.119 | 0.090 | -1.324 | 0.185 |
|  | **larger group size (scaled)** | **-0.334** | **0.114** | **-2.922** | **0.003** |
| Ecological | **Prior season type [dry]** | **1.934** | **0.272** | **7.118** | **<0.001** |
|  | Prior season type [wet] | -1.182 | 0.133 | 1.364 | 0.172 |

## Discussion

Overall, non-territorial groups of vulturine guineafowls expressed fidelity in their space use (consistent home ranges across time). Group home ranges overlapped with those of other groups, but these overlaps were substantially lower than home range fidelity expressed by groups, suggesting group home ranges are distinct from each other. In other words, groups did not randomly use the whole area available to them but preferred moving within a subset of the available area, and this area was distinct from most other groups. Even though the home range fidelity of groups was high, we also detected extensive temporal variation—even when making comparisons within the same set of seasonal conditions. We found that both social processes and the carryover effects from the recent ecological conditions predicted some of the temporal variations in the fidelity of preferred (intermediate season) home ranges of groups. Many of these spatial and social processes also affected the overlap in the home ranges between groups.

Home ranges, and home range overlap among groups, fundamentally impact how individuals experience their environment. The more consistent the environment is, the better groups can use prior experience to exploit resources (Riotte-Lambert & Matthiopoulos, 2020), but this ability depends on the ability of groups to maintain that experience among group members. At the same time, having more overlap with other groups increases the uncertainty arising from the foraging decisions of other groups (in terms of what resources they might have already exploited), with overlap being less preferable when resources are scarcer (i.e. the environment is drier). We found that the spatial and social processes that affect home range fidelity and overlap among groups aligned with these expectations.

We found that greater social change, longer periods between intermediate seasonal conditions, and drier conditions prior to the second sampling period reduced home range fidelity. Experiencing a greater turnover in memberships means that groups have more diverse sets of preferences (or information) among group members (Stanley et al., 2008). When groups express shared decision-making as vulturine guineafowl groups typically do (Papageorgiou, Nyaguthii, et al., 2024), having more diverse sets of preferences as a result of experiencing greater turnover in group membership could lead to different collective decisions (Chimento et al., 2021). This would result in changes in space use relative to earlier periods (and thus a lower home range fidelity). Vulturine guineafowl groups typically leave the area during drier periods (Papageorgiou et al., 2021). When groups do not use their home range for a long time, individuals may remember less about the area and the surrounding environment itself may change (e.g. landmark trees might have died or been destroyed by megaherbivores), leading to changes in space use relative to previous periods when groups were using the same area. These findings highlight also that social and ecological processes are unlikely to be acting in isolation (Webber et al., 2023).

Importantly, we found that ecological conditions had a stronger effect in shaping the overall home ranges in the subsequent intermediate seasons and that social effects were more important in determining the use of core areas. The overall home ranges during dry seasons can partially overlap with home ranges during intermediate seasons (the median overlap index was previously suggested to be approximately 0.55 (Papageorgiou et al., 2021)). Thus, the overall home range use during the prior dry season may significantly modify the overall home range use in the subsequent intermediate seasons. By contrast, groups show a substantially greater shift in their core area during dry seasons, thus the core area fidelity may not be affected by the prior ecological conditions as much. The social effects on core areas are likely to be affected more by wet season dynamics. This is because the core area can represent the preferred area by groups for breeding during wet seasons (Nyaguthii et al., 2025). Thus, individuals can have strong preferences for the core area and may have stronger needs to exhibit their preferences during intermediate seasons that follow wetter conditions.

Overlap between groups also largely followed expectations. We found that home range overlap in intermediate seasons that followed drier periods was typically lower. This suggests that groups are responding to the slow return of resources after rainfall events, which is likely because it takes time for the environment to respond (e.g. grasses to seed, and grasses are an important part of the diet of vulturine guineafowl (Sikenykeny et al. *in prep*)). This result also suggests that inter-seasonal carryover effects likely shape the competition experienced among groups, such that the same seasonal condition (e.g. the same NDVI value) corresponds to more or less food depending on the previous season type. We also found that sets of groups with greater social contacts (i.e. those that were previously in the same supergroup and those that roosted in similar locations) also had greater home range overlap. Of note, groups that were previously part of a supergroup (a combination of two or more groups in the vulturine guineafowl multilevel society) typically had higher core area overlap, with larger differences in group size decreased home range overlap among pairs of groups. The signal of competition among vulturine guineafowl groups—which show no outward territorial behaviour—was also detectable through larger groups expressing smaller core area overlap with other groups.

An important aspect of our study is that we investigated patterns of both core and broader space use by groups. Groups spend substantial amounts of time in core areas for key activities such as foraging, resting, and sheltering from heat and predators during daytime, while they use other areas in the overall home ranges for less time, for example, for commuting between foraging patches. The main differences between the overall home ranges and core areas in movement decisions may come from the potential differences in how strong preferences individuals may have for the area and what they use the area for (e.g. survival during drier periods versus reproduction during wetter periods). Thus, exploring the differences in fidelity in the overall home ranges and in the core areas highlights the importance of differentiating the biological functions of these areas. It also gives insights into segregation in space that arises due to key survival functions, i.e. competition for food resources, versus those that arise due to reproductive opportunities, i.e. competition for mates.

The current study gives insights into the emergence of territoriality. Groups showed the lowest overlap in core areas, especially when they recently experienced milder ecological conditions. This suggests that carryover from conditions suitable for breeding is potentially driving the partitioning of core areas. There may be competition for females, as we observe that males occasionally engage in mate defence from males in other groups during the wetter ecological conditions. Despite the increased expression of spatial partitioning among groups, vulturine guineafowl never establish territories. This is likely because the benefits would not outweigh the large costs involved (Brown, 1964; Mezzini et al., 2025)). For example, vulturine guineafowl forage on grasses and insects at large and open glades in savanna, which can be difficult to monopolise or defend from competitors. Yet, even without behaviours that actively exclude competing groups, vulturine guineafowl are still likely to experience competition and thus benefit from using different spaces to what other groups use. This study thus suggests that animals may still actively (consciously or not) partition space use even when there are no typical territorial behaviours, like aggressive defence of resources or space (e.g. Tórrez-Herrera et al., 2020).

Overall, this study provides insights into how social and ecological factors can shape the movement decisions by groups about where to go, captured as the temporal variations in consistent space use by non-territorial groups. High consistency in space use means that vulturine guineafowl populations are partitioning space across groups in a consistent way. Doing so may enable them to minimize competition, and we found several lines of evidence suggesting that space use is more clearly partitioned when conditions are less favourable. These patterns could also explain why vulturine guineafowl do not show a monotonic increase in social contact with increasing population density (Albery et al., 2024). The exact decision-making process that determines where groups range and the behavioural mechanisms for them to maintain the distinct space use even when groups share their key locations (e.g. roosting sites) remain to be investigated. Furthermore, this study gives insights into one pathway to emerge territorial-like space use among groups as they experience fluctuating levels of competition. Future research could use experimental manipulations of resources to test if non-territorial groups with highly overlapped home ranges can develop territorial-like space use in the field when the benefits of defending resource patches are modified to outweigh the costs. This can give further insights into how competition can facilitate the emergence of territoriality (Ogino & Farine, 2024; Riotte-Lambert et al., 2015) and how group-living can play a role in enhancing the process.

## Acknowledgements

We thank Charlotte Christensen, James Klarevas-Irby, and the Farine lab for their feedback and comments. We appreciate the help from the past and current members of the vulturine guineafowl fieldwork team, the Mpala Research Centre and the National Museums of Kenya for their support of the project.

## Author contributions

MO and DRF developed the project. All authors contributed to the data collection and scientific discussion. MO analyzed the data and wrote the first draft, and MO and DRF revised the manuscript.

## References

Albery, G. F., Becker, D. J., Firth, J. A., Silk, M., Sweeny, A. R., Wal, E. V., Webber, Q., Allen, B., Babayan, S. A., & Barve, S. (2024). Density-dependent network structuring within and across wild animal systems. BioRxiv, 2024.2006. 2028.601262.

Andersson, M. (1978). Optimal foraging area: size and allocation of search effort. Theoretical population biology, 13(3), 397–409.

Andersson, M. (1981). Central place foraging in the whinchat, Saxicola rubetra. Ecology, 62(3), 538–544.

Beauchamp, G., & Ruxton, G. D. (2005). Harvesting resources in groups or alone: the case of renewing patches. Behavioral Ecology, 16(6), 989–993.

Brooks, M. E., Kristensen, K., Van Benthem, K. J., Magnusson, A., Berg, C. W., Nielsen, A., Skaug, H. J., Mächler, M., & Bolker, B. M. (2017). glmmTMB balances speed and flexibility among packages for zero-inflated generalized linear mixed modeling.

Brown, C. R., Brown, M. B., & Brazeal, K. R. (2008). Familiarity with breeding habitat improves daily survival in colonial cliff swallows. Animal Behaviour, 76(4), 1201–1210.

Brown, J. L. (1964). The evolution of diversity in avian territorial systems. The Wilson Bulletin, 160–169.

Burt, W. H. (1943). Territoriality and home range concepts as applied to mammals. Journal of mammalogy, 24(3), 346–352.

Calabrese, J. M., Fleming, C. H., & Gurarie, E. (2016). ctmm: An R package for analyzing animal relocation data as a continuouslJtime stochastic process. Methods in Ecology and Evolution, 7(9), 1124–1132.

Chimento, M., Alarcón-Nieto, G., & Aplin, L. M. (2021). Population turnover facilitates cultural selection for efficiency in birds. Current Biology, 31(11), 2477-2483. e2473.

Christensen, C., & Radford, A. N. (2018). Dear enemies or nasty neighbors? Causes and consequences of variation in the responses of group-living species to territorial intrusions. Behavioral Ecology, 29(5), 1004–1013.

Clark, C. W., & Mangel, M. (1984). Foraging and flocking strategies: information in an uncertain environment. The American Naturalist, 123(5), 626–641.

Clarke, M. F., Da Silva, K. B., Lair, H., Pocklington, R., Kramer, D. L., & McLaughlin, R. L. (1993). Site familiarity affects escape behaviour of the eastern chipmunk, Tamias striatus. Oikos, 533–537.

Conradt, L. (2012). Models in animal collective decision-making: information uncertainty and conflicting preferences. Interface focus, 2(2), 226–240.

Davis, G. H., Crofoot, M. C., & Farine, D. R. (2022). Using optimal foraging theory to infer how groups make collective decisions. Trends in Ecology & Evolution, 37(11), 942–952.

Ekman, J., Cederholm, G., & Askenmo, C. (1981). Spacing and survival in winter groups of Willow Tit Parus montanus and Crested Tit P. cristatus--a removal study. The Journal of Animal Ecology, 1–9.

Elliott, K. H., Woo, K. J., Gaston, A. J., Benvenuti, S., Dall’Antonia, L., & Davoren, G. K. (2009). Central-place foraging in an Arctic seabird provides evidence for Storer-Ashmole’s halo. The Auk, 126(3), 613–625.

Fagan, W. F., Lewis, M. A., AugerlJMéthé, M., Avgar, T., Benhamou, S., Breed, G., LaDage, L., Schlägel, U. E., Tang, W. w., & Papastamatiou, Y. P. (2013). Spatial memory and animal movement. Ecology letters, 16(10), 1316–1329.

Fishlock, V., Caldwell, C., & Lee, P. C. (2016). Elephant resource-use traditions. Animal Cognition, 19, 429–433.

Fleming, C. C., Justin. (2022). ctmm: Continuous-Time Movement Modeling, version 1.1.0. R Package. https://CRAN.R-project.org/package=ctmm

Fretwell, S. D., & Calver, J. S. (1969). On territorial behavior and other factors influencing habitat distribution in birds: II. Sex ratio variation in the Dickcissel (Spiza americana Gmel). Acta biotheoretica, 19(1), 37–44.

Galef Jr, B. G., & Whiskin, E. E. (1997). Effects of social and asocial learning on longevity of food-preference traditions. Animal Behaviour, 53(6), 1313–1322.

Gallagher, A. J., Creel, S., Wilson, R. P., & Cooke, S. J. (2017). Energy landscapes and the landscape of fear. Trends in Ecology & Evolution, 32(2), 88–96.

Gaynor, K. M., Brown, J. S., Middleton, A. D., Power, M. E., & Brashares, J. S. (2019). Landscapes of fear: spatial patterns of risk perception and response. Trends in Ecology & Evolution, 34(4), 355–368.

Grist, H., Daunt, F., Wanless, S., Nelson, E. J., Harris, M. P., Newell, M., Burthe, S., & Reid, J. M. (2014). Site fidelity and individual variation in winter location in partially migratory European shags. PLoS One, 9(6), e98562.

Grueter, C. C., Qi, X., Zinner, D., Bergman, T., Li, M., Xiang, Z., Zhu, P., Migliano, A. B., Miller, A., & Krützen, M. (2020). Multilevel organisation of animal sociality. Trends in Ecology & Evolution, 35(9), 834–847.

Harel, R., Loftus, J. C., & Crofoot, M. C. (2021). Locomotor compromises maintain group cohesion in baboon troops on the move. Proceedings of the Royal Society B, 288(1955), 20210839.

Hartig, F. (2022). DHARMa: residual diagnostics for hierarchical (multi-level/mixed) regression models. R package version 0.4. 6. In.

Hendrix, J., Robitaille, A., Kusch, J., Webber, Q., & Vander Wal, E. (2024). Faithful pals and familiar locales: differentiating social and spatial site fidelity during reproduction. Philosophical Transactions B, 379(1912), 20220525.

Klarevas-Irby, J. A., Nyaguthii, B., & Farine, D. R. (2025). Moving as a group imposes constraints on the energetic efficiency of movement. Proceedings B, 292(2041), 20242760.

Krebs, J. R. (1971). Territory and breeding density in the Great Tit, Parus major L. Ecology, 52(1), 2–22.

Madsen, A. E., Lyon, B. E., Chaine, A. S., Block, T. A., & Shizuka, D. (2023). Loss of flockmates weakens winter site fidelity in golden-crowned sparrows (Zonotrichia atricapilla). Proceedings of the National Academy of Sciences, 120(32), e2219939120.

Mezzini, S., Fleming, C. H., Medici, E. P., & Noonan, M. J. (2025). How resource abundance and resource stochasticity affect organisms’ range sizes. Movement ecology, 13(1), 1–14.

Mirville, M. O., Ridley, A. R., Samedi, J., Vecellio, V., Ndagijimana, F., Stoinski, T. S., & Grueter, C. C. (2020). Intragroup behavioral changes following intergroup conflict in mountain gorillas (Gorilla beringei beringei). International Journal of Primatology, 41(2), 382–400.

Nyaguthii, B., Dehnen, T., KlarevaslJIrby, J. A., Papageorgiou, D., Kosgey, J., & Farine, D. R. (2025). Cooperative and plural breeding by the precocial Vulturine Guineafowl. Ibis.

Ogino, M., & Farine, D. R. (2024). Collective intelligence facilitates emergent resource partitioning through frequency-dependent learning. Philosophical Transactions B, 379(1909), 20230177.

Ogino, M., Strauss, E. D., & Farine, D. R. (2023). Challenges of mismatching timescales in longitudinal studies of collective behaviour. Philosophical Transactions of the Royal Society B, 378(1874), 20220064.

Ogle, D. D., Jason Wheeler, Powell; Dinno, Alexis. (2025). FSA: Simple Fisheries Stock Assessment Methods. R package version 0.9.6. https://CRAN.R-project.org/package=FSA

Papageorgiou, D., Cherono, W., Gall, G., Nyaguthii, B., & Farine, D. R. (2024). Testing the information centre hypothesis in a multilevel society. Journal of Animal Ecology, 93(8), 1147–1159.

Papageorgiou, D., Christensen, C., Gall, G. E., Klarevas-Irby, J. A., Nyaguthii, B., Couzin, I. D., & Farine, D. R. (2019). The multilevel society of a small-brained bird. Current Biology, 29(21), R1120–R1121.

Papageorgiou, D., & Farine, D. R. (2020). Group size and composition influence collective movement in a highly social terrestrial bird. Elife, 9, e59902.

Papageorgiou, D., & Farine, D. R. (2021). Multilevel societies in birds. Trends in Ecology & Evolution, 36(1), 15–17.

Papageorgiou, D., Nyaguthii, B., & Farine, D. R. (2024). Compromise or choose: shared movement decisions in wild vulturine guineafowl. Communications Biology, 7(1), 95.

Papageorgiou, D., Rozen-Rechels, D., Nyaguthii, B., & Farine, D. R. (2021). Seasonality impacts collective movements in a wild group-living bird. Movement ecology, 9, 1–12.

Piper, W. H. (2011). Making habitat selection more “familiar”: a review. Behavioral Ecology and Sociobiology, 65, 1329–1351.

Ranc, N., Moorcroft, P. R., Ossi, F., & Cagnacci, F. (2021). Experimental evidence of memory-based foraging decisions in a large wild mammal. Proceedings of the National Academy of Sciences, 118(15), e2014856118.

Riotte-Lambert, L., Benhamou, S., & Chamaillé-Jammes, S. (2015). How memory-based movement leads to nonterritorial spatial segregation. The American Naturalist, 185(4), E103–E116.

Riotte-Lambert, L., & Matthiopoulos, J. (2020). Environmental predictability as a cause and consequence of animal movement. Trends in Ecology & Evolution, 35(2), 163–174.

Rohwer, F. C., & Anderson, M. G. (1988). Female-biased philopatry, monogamy, and the timing of pair formation in migratory waterfowl. In Current ornithology (pp. 187–221). Springer.

Shepard, E. L., Wilson, R. P., Rees, W. G., Grundy, E., Lambertucci, S. A., & Vosper, S. B. (2013). Energy landscapes shape animal movement ecology. The American Naturalist, 182(3), 298–312.

Somershoe, S. G., Brown, C. D., & Poole, R. T. (2009). Winter site fidelity and over-winter site persistence of passerines in Florida. The Wilson Journal of Ornithology, 121(1), 119–125.

Stanley, E. L., Kendal, R. L., Kendal, J. R., Grounds, S., & Laland, K. N. (2008). The effects of group size, rate of turnover and disruption to demonstration on the stability of foraging traditions in fish. Animal Behaviour, 75(2), 565–572.

Tanner, J. C., & Hemingway, C. T. (2025). Choice overload and its consequences for animal decision-making. Trends in Cognitive Sciences.

Team, R. C. (2021). R: A language and environment for statistical computing. R Foundation for Statistical Computing, Vienna, Austria. https://www.R-project.org/.

Tórrez-Herrera, L. L., Davis, G. H., & Crofoot, M. C. (2020). Do monkeys avoid areas of home range overlap because they are dangerous? A test of the risk hypothesis in white-faced capuchin monkeys (Cebus capucinus). International Journal of Primatology, 41, 246–264.

van der Post, D. J., & Semmann, D. (2011). Patch depletion, niche structuring and the evolution of co-operative foraging. BMC Evolutionary Biology, 11, 1–17.

Webber, Q. M., Albery, G. F., Farine, D. R., PinterlJWollman, N., Sharma, N., Spiegel, O., Vander Wal, E., & Manlove, K. (2023). Behavioural ecology at the spatial–social interface. Biological Reviews, 98(3), 868–886.

Wilson, E. O. (2000). Sociobiology: The new synthesis. Harvard University Press.

Winner, K., Noonan, M. J., Fleming, C. H., Olson, K. A., Mueller, T., Sheldon, D., & Calabrese, J. M. (2018). Statistical inference for home range overlap. Methods in Ecology and Evolution, 9(7), 1679–1691.

Yoder, J. M., Marschall, E. A., & Swanson, D. A. (2004). The cost of dispersal: predation as a function of movement and site familiarity in ruffed grouse. Behavioral Ecology, 15(3), 469–476.

